# Oxidative Phosphorylation (OXPHOS) Promotes the Formation and Growth of Melanoma Lung and Brain Metastases

**DOI:** 10.1101/2025.01.23.633049

**Authors:** Renato A. Guerrieri, Grant M. Fischer, David A. Kircher, Aron Y. Joon, Jacob R. Cortez, Allie H. Grossman, Courtney W. Hudgens, Debora A Ledesma, Rossana Lazcano, Christian YB Onana, Barbara G. Knighton, Swaminathan Kumar, Qianghua Hu, Y. N. Vashisht Gopal, Jennifer L. McQuade, Wanleng Deng, Lauren E. Haydu, Jeffrey E. Gershenwald, Alexander J. Lazar, Michael T. Tetzlaff, Sheri L. Holmen, Michael A. Davies

**Author notes:** Co-First Authors (Renato A. Guerrieri, Grant M. Fischer, David A. Kircher). Co-Senior Authors (Sheri L. Holmen, Michael A. Davies). Corresponding Author: Michael A. Davies, 1515 Holcombe Blvd., Unit 0430, Houston, TX 77030.

## Abstract

Melanoma mortality is driven by the formation and growth of distant metastases. Here, we interrogated the role of tumor oxidative phosphorylation (OXPHOS) in the formation of distant metastases in melanoma. OXPHOS was the most upregulated metabolic pathway in primary tumors that formed distant metastases in the RCAS-TVA mouse model of spontaneous lung and brain metastases, and in melanoma patients that developed brain or other distant metastases. Knockout of PGC1α in melanocytes in the RCAS-TVA melanoma mouse model had no impact on primary tumor formation, but markedly reduced the incidence of lung and brain metastases. Genetic knockout of a component of electron transport chain complex I, NDUFS4, in B16-F10 and D4M-UV2 murine melanoma cell lines did not impact tumor incidence following subcutaneous, intravenous, or intracranial injection, but decreased tumor burden specifically in the lungs and brain. Together, these data demonstrate that OXPHOS is critical for the formation of metastases in melanoma.

**STRUCTURED ABSTRACT:** *Purpose:* Melanoma mortality is driven by the formation and growth of distant metastases. However, the process and pathogenesis of melanoma metastasis remain poorly understood. Here, we interrogate the role of tumor oxidative phosphorylation (OXPHOS) in the formation of distant metastases in melanoma.

*Experimental Design:* This study includes (1) new RNA-seq analysis of primary melanomas from patients characterized for distant metastasis events; (2) RNA-seq analysis and functional testing of genetic OXPHOS inhibition (PGC1α KO) the RCAS-TVA model, which is the only existing immunocompetent murine model of autochthonous lung and brain metastasis formation from primary melanoma tumors; and (3) functional experiments of genetic OXPHOS inhibition (NDUFS4 KO) in the B16-F10 and D4M-UV2 murine melanoma cell lines, including evaluation of subcutaneous, lung, and brain metastatic site dependencies.

*Results:* OXPHOS was the most upregulated metabolic pathway in primary tumors that formed distant metastases in the RCAS-TVA mouse model of spontaneous lung and brain metastases, and in melanoma patients that developed brain or other distant metastases. Knockout of PGC1a in melanocytes in the RCAS-TVA melanoma mouse model had no impact on primary tumor formation, but markedly reduced the incidence of lung and brain metastases. Genetic knockout of a component of electron transport chain complex I, NDUFS4, in B16-F10 and D4M-UV2 murine melanoma cell lines did not impact tumor incidence following subcutaneous, intravenous, or intracranial injection, but decreased tumor burden specifically in the lungs and brain.

*Conclusions:* Together, these data demonstrate that OXPHOS is critical for the formation of metastases in melanoma.

**TRANSLATIONAL RELEVANCE:** Melanoma is the most aggressive form of skin cancer. One hallmark of this disease is a high risk of distant metastasis formation. The process and pathogenesis of metastasis in this disease remain poorly understood and there is controversy regarding the role of oxidative phosphorylation (OXPHOS) in melanoma metastasis. This study incorporates RNAseq analysis of primary melanoma tumors from patients characterized for distant metastasis events, RNAseq analysis of the only existing immunocompetent murine model of autochthonous lung and brain metastasis formation from primary melanoma tumors, and functional testing in multiple syngeneic models of melanoma at different tissue sites. This integrated analysis consistently demonstrates that melanoma OXPHOS promotes distant metastasis to the lungs and brain, two of the most common and clinically relevant sites of melanoma metastasis. This improved understanding of tumor OXPHOS may represent novel vulnerabilities for therapeutics development and surveillance/preventative strategies for melanoma metastasis.

## INTRODUCTION

Melanoma is among the deadliest forms of skin cancer, accounting for over 80% of skin cancer-related deaths.^[1, 2]^ Mortality from melanoma results from metastasis of primary tumors to distant organ sites, which occurs more frequently and earlier in disease progression in melanoma compared to most other solid tumors.^[3, 4]^ Melanoma brain metastasis (MBM) is a particularly devastating complication of this disease that drives morbidity and mortality.^[5]^ Despite markedly improved outcomes of melanoma patients due to advances in immune and targeted therapies, the median overall survival from MBM remains approximately one year.^[6–9]^ Improved understanding of the factors that predict and promote MBM formation could lead to rational treatment and/or prevention strategies to reduce their incidence and morbidity.

Previously, we found that MBMs are significantly enriched in OXPHOS gene expression compared to extracranial metastases (ECMs).^[10]^ In addition, treatment of melanomas generated in the RCAS-TVA mouse model with IACS-010759, a potent OXPHOS inhibitor targeting electron transport chain complex I, showed OXPHOS inhibition completely inhibited or eradicated MBMs, but had no impact on primary tumor growth or lung metastases.^[10, 11]^ Together these results strongly implicate OXPHOS in the formation of MBMs, a finding which was subsequently confirmed in brain metastases from other tumor types.^[12–15]^ In addition, we found that MBMs with elevated OXPHOS are associated with poorer survival outcomes.^[16]^ However, the role of OXPHOS as a risk factor for MBM, and functionally as a mediator of MBM formation, remains unclear. These are important questions to address as other studies have suggested that oxidative stress and PGC1α, which can be associated with OXPHOS, inhibits distant metastasis formation.^[17–19]^

To address this knowledge gap, we analyzed the role of OXPHOS in MBM formation. These integrated studies of primary tumors (from mice and melanoma patients) and melanoma metastasis models strongly implicate OXPHOS in lung and brain metastasis in this disease.

## RESULTS

### Oxidative Phosphorylation in primary tumors from a murine model of melanoma is associated with risk of lung and brain metastasis

To determine the role of tumor OXPHOS in the formation of melanoma metastases, we first analyzed primary tumors from the *Dct::TVA;BRAF^CA^;Cdkn2a^lox/lox^; Pten^lox/lox^* (BCP) RCAS-TVA mouse model. These mice develop primary tumors and spontaneous lung and brain metastases when subcutaneously injected with viruses encoding mutant AKT in combination with RCAS-Cre. Delivery of RCAS-Cre results in expression of *BRAF*^V600E^ and genetic loss of *Cdkn2a* and *Pten*.^[11]^ For mice injected with viruses encoding Cre and E17K activating mutants of AKT1, AKT2, and AKT3, brain metastases were detected in 38%, 19%, and 11% of mice and lung metastases were detected in 24%, 19%, and 21% of mice, respectively.^[20]^ Delivery of RCAS-Cre alone to these mice resulted in no brain metastases and lung metastases in only 8% of mice. We performed RNAseq on a subset of primary tumors from these mice (**Table S1**) and used pre-ranked gene set enrichment (GSEA-P) analysis to compare metabolic pathway expression (**Table S2**) between primary tumors from the mice that developed metastases to the brain and/or lungs (“P_AllAkt_Met”) compared to mice that did not (“P_AllAkt_NoMet”). The KEGG_OXPHOS gene set was the most significantly enriched metabolic pathway (FDR *q*-val <0.05) in primary tumors from P_AllAkt_Met mice compared to P_AllAkt_NoMet mice (**Figure 1A**). Consistent with this association, primary tumor OXPHOS-Index (OP-Index^[10]^) scores were significantly elevated in P_AllAkt_Met mice compared to P_AllAkt_NoMet mice (*p*=0.0008) (**Figure 1B**). Similar results were observed in the comparison of P_AllAkt_Met mice with P_WTAkt_NoMet mice (**Figure 1C-D**).

**Figure 1:**
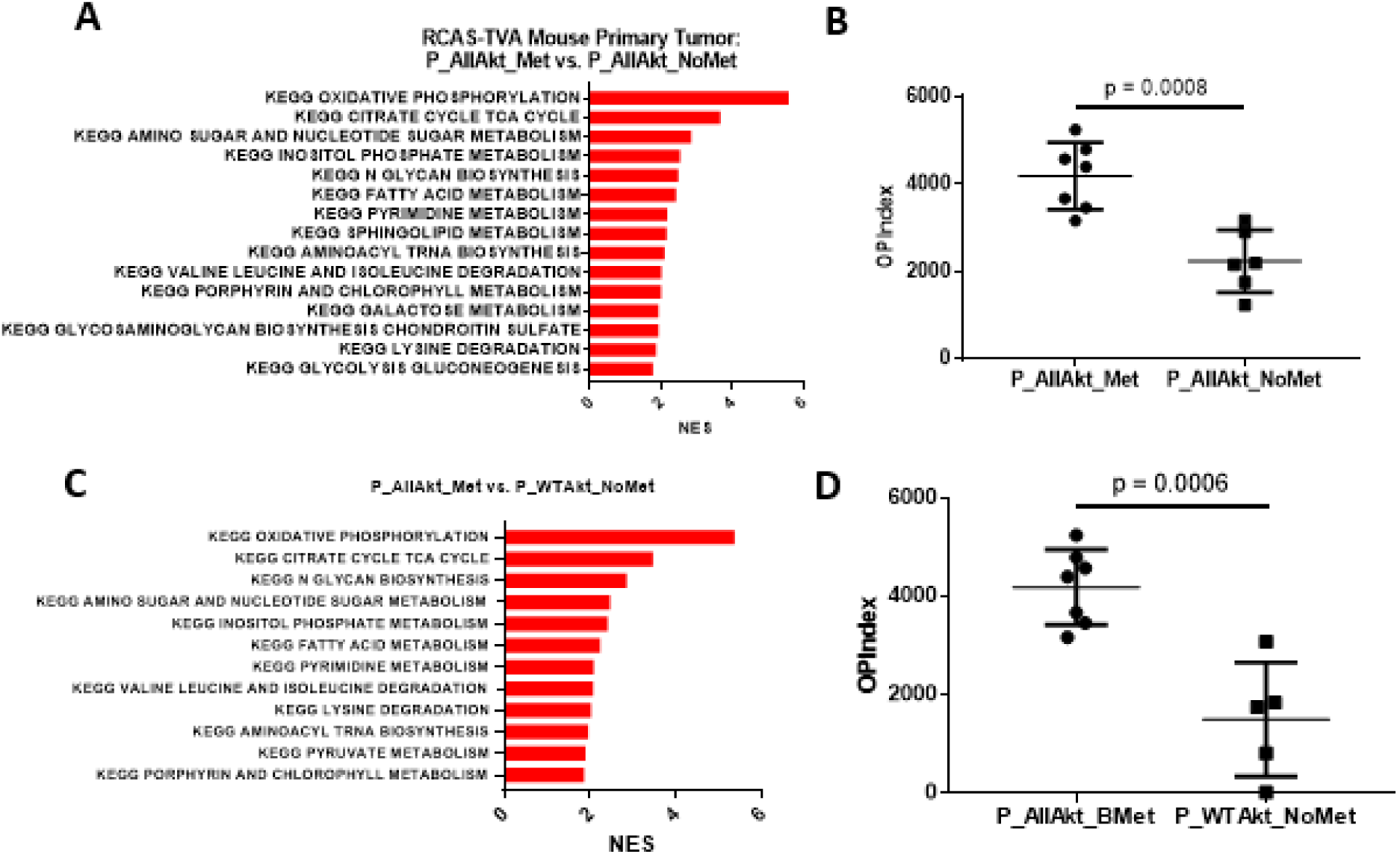
Oxidative phosphorylation associates with lung and brain metastasis formation from primary tumors from an RCAS-TVA model of autochthonous melanoma metastasis. **(A)** GSEA-P analysis enrichment plot demonstrating all KEGG metabolism pathways significantly enriched (FDR q-val<0.05) in BCP ± AKT1/2/3-E17K metastatic primaries vs. BCP ± AKT1/2/3-E17K non-metastatic primaries. Normalized enrichment score (NES) forms the x-axis. **(B)** OP-Indices of BCP ± AKT1/2/3-E17K metastatic primaries and BCP ± AKT1/2/3-E17K non-metastatic primaries. Each dot represents a single sample and lines indicate mean +/- S.D. Significance determined via two-sided Student’s *t*-test. **(C)** GSEA-P analysis enrichment plot demonstrating all KEGG metabolism pathways significantly enriched (FDR q-val<0.05) in BCP ± AKT1/2/3-E17K metastatic primaries vs. BCP without AKT1/2/3-E17K non-metastatic primaries. Normalized enrichment score (NES) forms the x-axis. **(D)** OP-Indices of BCP ± AKT1/2/3-E17K metastatic and BCP without AKT1/2/3-E17K non-metastatic primaries. Each dot represents a single sample and lines indicate mean +/- S.D. Significance determined via two-sided Student’s *t*-test.

### Oxidative Phosphorylation in patient primary tumors is associated with incidence of melanoma metastasis

We next evaluated the association of upregulated OXPHOS with metastatic risk in melanoma patients. Building upon a previously reported RNAseq analysis of archival FFPE primary melanoma specimens,^[21]^ we expanded the cohort to include additional primary tumors to allow for assessment of associations with MBM risk. The final cohort included primary tumors from melanoma patients who developed MBMs within 3-18 months of initial diagnosis (“P_MBM”; n=19); extracranial metastases (ECMs) only to non-CNS sites within 3-18 months (“P_ECM”; n=16); and no distant metastases within 60 months (“P_NoMet”; n=19). GSEA-P analysis found significant enrichment in KEGG OXPHOS gene set expression in primary melanomas that developed metastases (“P_AllMet” = P_MBM+P_ECM) compared to those that did not (P_NoMet; FDR *q*-val=0.002) (**Figure 2A-B**). P_MBM and P_ECM tumors each showed elevated KEGG OXPHOS gene set expression compared to P_NoMet (FDR *q*-val=0.048 and <0.001, respectively) (**Figures 2C-D**). However, there was no significant difference in KEGG OXPHOS gene set expression between P_MBM and P_ECM tumors (FDR *q*-val=0.516) (**Figure 2E**). Since OXPHOS can be associated with oxidative stress, we performed GSEA-P for a curated pathway set, which indicated that several oxidative stress pathways were significantly enriched in P_AllAkt_Met versus P_AllAkt_NoMet murine primary tumors, and in P_AllMet versus P_NoMet patient samples (**Table S3-S4**).

**Figure 2:**
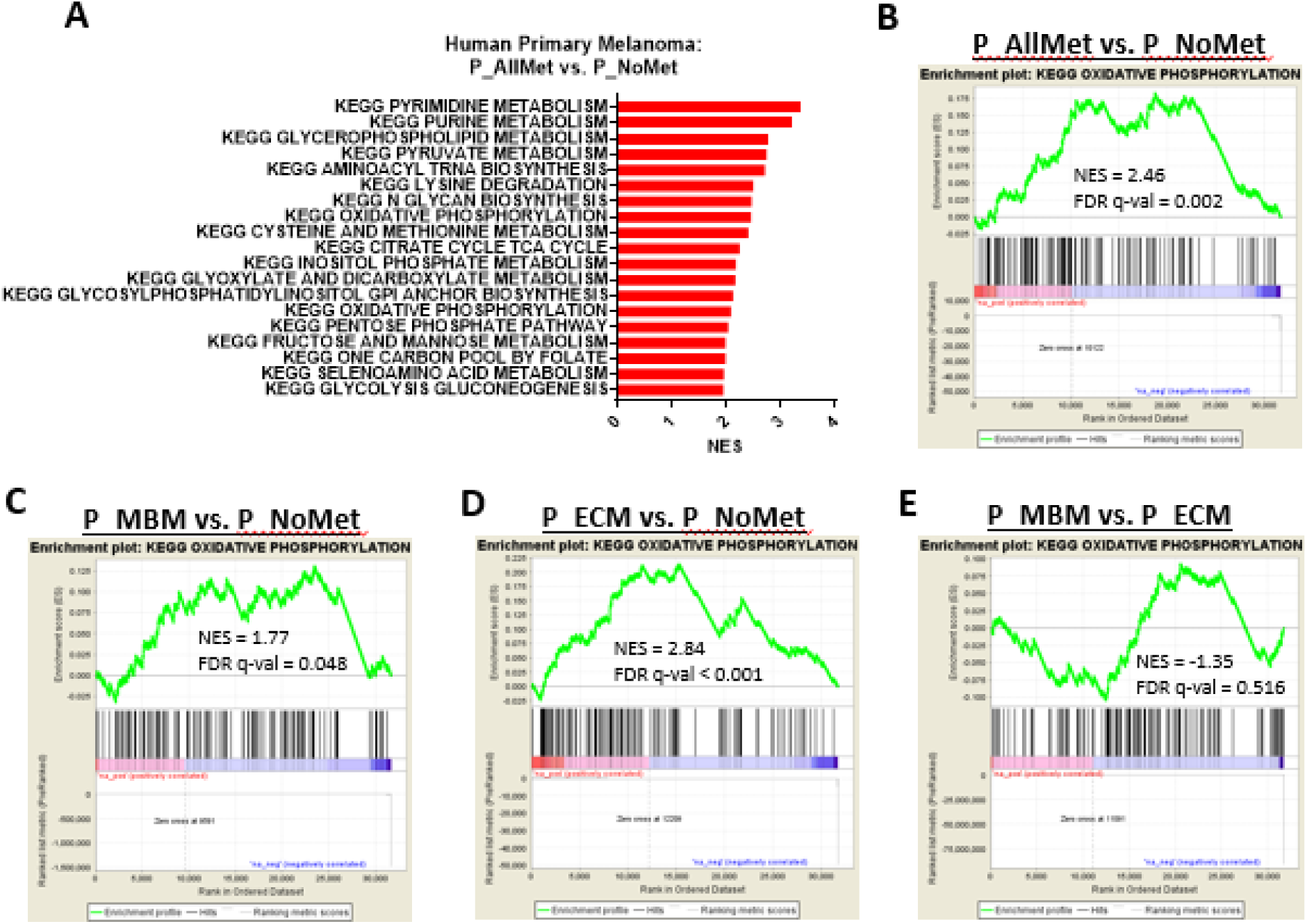
Oxidative phosphorylation associates with metastasis formation from primary tumors from human primary melanomas. **(A)** GSEA-P analysis enrichment plot demonstrating all KEGG metabolism pathways significantly enriched (FDR q-val<0.05) in P_AllMet metastatic primaries vs. P_NoMet non-metastatic primaries. Normalized enrichment score (NES) forms the x-axis. **(B-D)** GSEA-P analysis demonstrating significant enrichment of the KEGG OXPHOS gene set in metastatic human primary melanomas (P_AllMet; n=35) **(B)**, brain-metastatic human primary melanomas (P_MBM; n=19) **(C)**, extracranially-only metastatic human primary melanomas (P_ECM; n=16) **(D)** each vs. non-metastatic human primary melanomas (P_NoMet; n=19) from 54 unique patients. **(E)** GSEA-P analysis demonstrating no significant enrichment of the KEGG OXPHOS gene set in P_MBM vs. P_ECM. NES and FDR q-val are listed on all enrichment plots.

We also tested whether OXPHOS in ECMs was associated with MBMs risk. We analyzed publicly available Illumina microarray data (GSE60464) from ECMs of stage IV melanoma patients who developed MBMs within 6 months of stage IV diagnosis (“MBM_Yes”; n=17) or did not develop MBMs within 18 months of stage IV diagnosis (“MBM_No”; n=25).^[22]^ There was no enrichment of KEGG OXPHOS or Hallmark OXPHOS gene sets between cerebrotropic and non-cerebrotropic ECMs (FDR *q*-val=0.917 and 0.856, respectively) (**Figure S1A-B**). Together, elevated OXPHOS in primary melanomas but not ECMs is associated with a higher risk of distant metastasis; moreover, this association is not specific for MBM formation.

### Functional knockout of PGC1α reduces incidence of lung and brain metastases in murine model of autochthonous melanoma metastasis

To investigate the impact of inhibiting OXPHOS at primary tumor initiation on melanoma metastasis formation, we utilized mice carrying a conditional (floxed) allele of peroxisome proliferative activated receptor, gamma, coactivator 1 alpha (*Ppargc1a; Pgc1a*). LoxP sites flank exons 3-5 of the *Ppargc1a* gene, which encodes for PGC1α, a key regulator of energy metabolism.^[23]^ *Pgc1α^lox/lox^*mice were crossed with RCAS-TVA BCP mice to generate *Dct::TVA;BRAF^CA^;Cdkn2a^lox/lox^:Pten^lox/lox^*;*Pgc1α^lox/lox^* (BCPP) mice. Newborn BCP and BCPP mice were subcutaneously injected with viruses encoding either RCAS-Cre alone or RCAS-Cre in combination with myristoylated (myr) AKT1 and monitored for tumor development (**Figure S2A**). Mice were evaluated for the presence of metastases post-mortem.

All injected mice developed primary tumors, indicating that genetic inhibition of PGC1α, and thus of OXPHOS, had no impact on tumor initiation. However, loss of PGC1α resulted in a significant improvement in overall survival in BCPP + myrAKT1 mice compared to BCP + myrAKT1 mice (*p*<0.0001) (**Figure 3A**). The overall survival for the BCP and BCPP RCAS-TVA mice was driven by a combination of both primary and metastatic tumor growth. All mice were processed and histologically examined for metastases to the lungs and brains; 71% of BCP + myrAKT1 mice developed lung metastases, yet the incidence of lung metastases was only 11% in BCPP + myrAKT1 mice (*p*=0.0006) (**Figure 3B**). Similarly, 79% of BCP + myrAKT1 mice developed brain metastases compared to only 5% of the BCPP + myrAKT1 mice (*p*<0.0001) (**Figure 3C**). The majority of BCP + myrAKT1 mice were sacrificed due to neurologic symptoms or respiratory distress, which is consistent with brain or lung metastases; mice in all other groups were sacrificed due to primary tumor burden and/or ulceration. PCR analysis confirmed loss of the floxed PGC1α allele in primary tumors of BCPP mice except for one mouse with lung metastases and one mouse with lung and brain metastases in the BCPP + myrAKT1 cohort (**Figure S2B**). Thus, it is possible that metastases formed in these mice due to incomplete loss of PGC1α expression. Together, inhibiting melanoma PGC1α/OXPHOS at the earliest stages of tumor initiation prevented lung and brain metastasis formation but did not impact primary tumor formation or growth.

**Figure 3:**
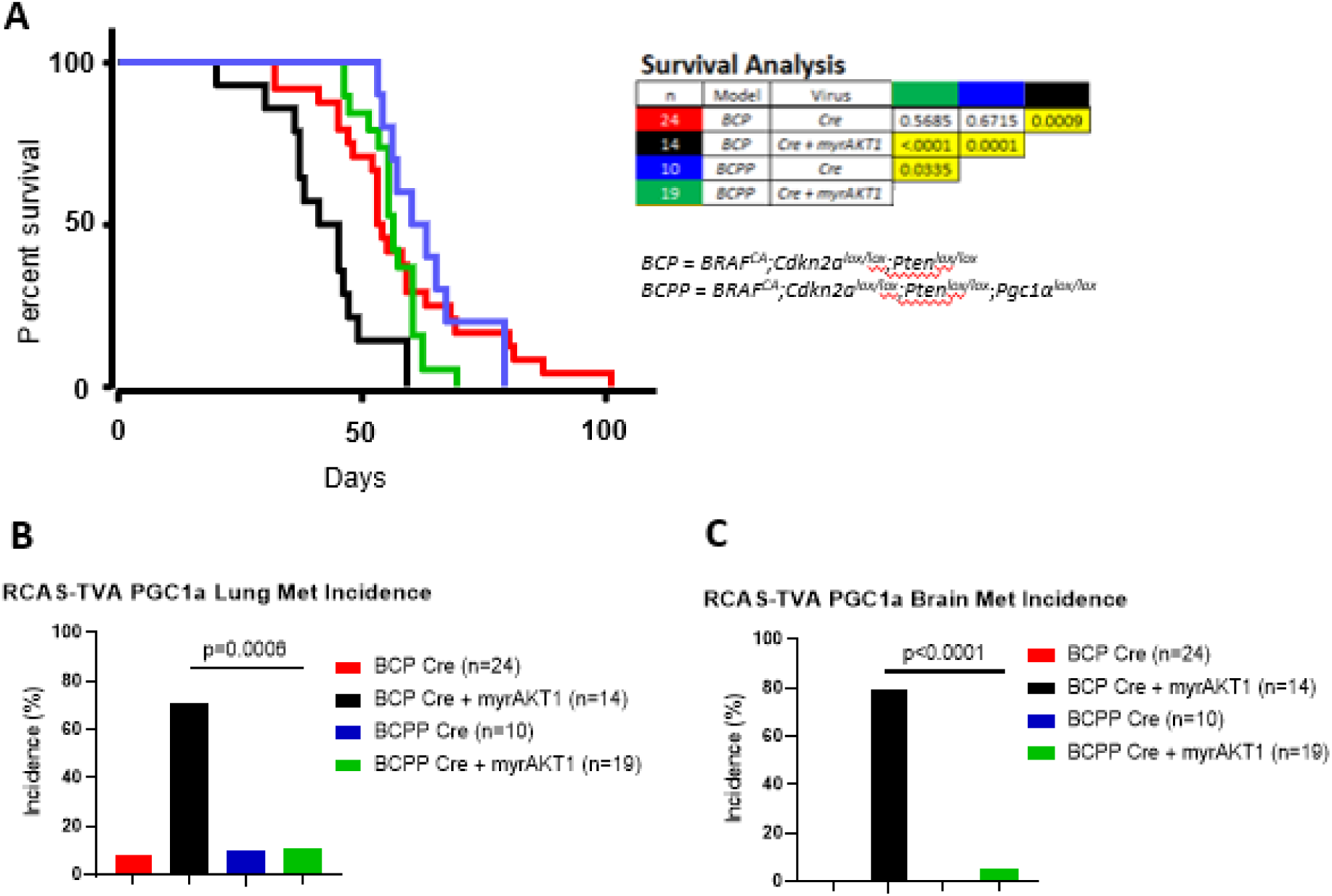
Loss of PGC1α/OXPHOS reduces metastasis and increases survival in an RCAS-TVA model of spontaneous murine melanoma brain and lung metastasis. **(A)** Kaplan-Meier overall survival analysis in BCP ± myrAKT1 vs. BCPP ± myrAKT1 RCAS-TVA mice and accompanying Legend Table. “n” refers to number of mice and color denotes animal groups. “Model” refers to genotype of RCAS-TVA mice, genotypes listed in detail below. “Virus” refers to addition of Cre + myrAKT1 or Cre alone. Colored columns refer to the p-values between their respective animal groups, with significance determined by Log-rank (Mantel-Cox) test. **(B-C)** Incidence of lung **(B)** and brain **(C)** metastases in BCP Cre ± myrAKT1 vs. BCPP Cre ± myrAKT1. Y-axis indicates tumor incidence. Significance determined via Fisher’s exact test.

### Oxidative Phosphorylation inhibition suppresses tumor growth in lung and brain tumors but not subcutaneous tumors in syngeneic B16-F10 and D4M-UV2 models of melanoma

To further test the role of OXPHOS in melanoma lung and brain metastasis pathogenesis, we established immunocompetent experimental models using luciferase-tagged B16-F10 and D4M-UV2 melanoma cell lines. To directly target OXPHOS, we generated stable CRISPR knockout clones of mitochondrial complex I subunit, NADH:ubiquinone oxidoreductase subunit S4 (*Ndufs4*, NDUFS4 KO). NDUFS4 KO cells exhibit reduced OXPHOS, without affecting glycolysis or oxidative stress compared to Wildtype (WT) cells (**Figure S3-S4**).

To assess the impact of NDUFS4/OXPHOS inhibition on tumor formation and growth at different organ sites, we implanted WT and NDUFS4 KO B16-F10 and D4M-UV2 tumors subcutaneously, intravenously (via tail vein injection), and intracranially. To quantify tumor burden after intravenous or intracranial injection, we harvested a subset of mice and measured *ex vivo* bioluminescence.

NDUFS4 KO had no effect on tumor growth or survival in mice following subcutaneous tumor cell injection compared to mice injected with WT B16-F10 or D4M-UV2 cells (**Figures 4A-B; S5A-B)**. However, NDUFS4 KO resulted in lower lung tumor burden and improved survival following tail vein injection in both B16-F10 (*p*=0.0002 and *p*=0.0163, respectively) (**Figures 4C-D**) and D4M-UV2 (*p*=0.0007 and *p*=0.0017, respectively) (**Figures S5C-D**). Similarly, direct intracranial injection of NDUFS4 KO resulted in lower brain tumor growth and improved overall survival in both B16-F10 (*p*=0.0135 and *p*=0.0071, respectively) (**Figures 4E-F**) and D4M-UV2 (*p*=0.0049 and *p*=0.0023, respectively) (**Figures S5E-F**). IHC was performed on B16-F10 tumors in each group to determine whether changes in tumor burden were associated with changes in apoptosis (cleaved caspase 3; CC3), proliferation (Ki-67), or angiogenesis (VEGF) (**Figure 5**). Proliferation and angiogenesis markers showed no significant differences, but NDUFS4 KO resulted in increased CC3 in both lung metastases (*p*=0.0001) and intracranial tumors (*p*=0.0109) compared to B16-F10 WT, which is consistent with increased apoptosis. Collectively, these data indicate that NDUFS4/OXPHOS inhibition impairs tumor growth in the lungs and brain, but does not impact subcutaneous tumor growth.

**Figure 4:**
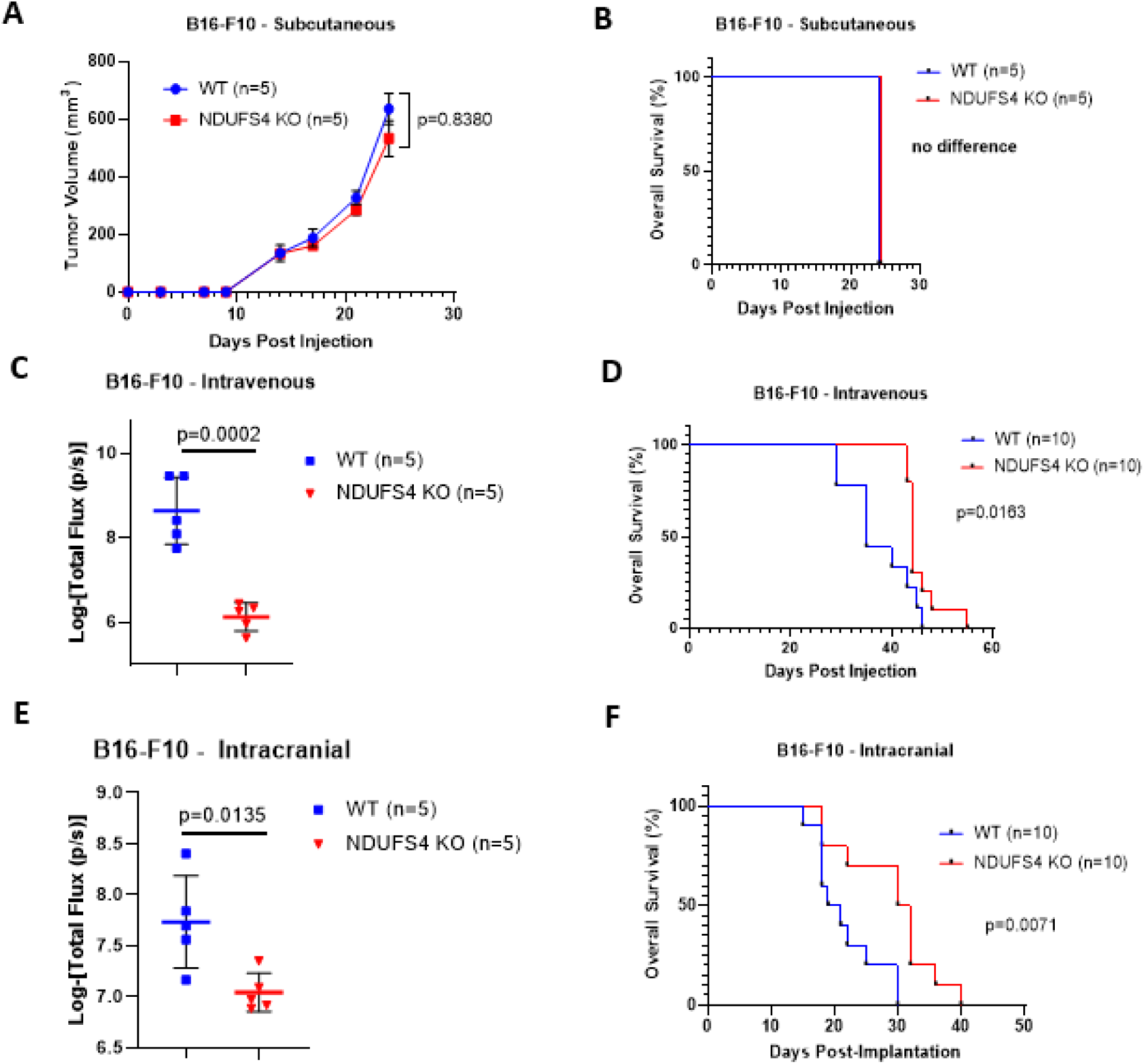
NDUFS4/OXPHOS is functionally significant for B16-F10 melanoma tumor growth and overall survival in syngeneic injection models of lung and brain, but not subcutaneous, tumor growth. **(A)** Growth of subcutaneous tumors. **(B, D, F)** Kaplan-Meier analysis of overall survival for mice bearing subcutaneous **(B)**, intravenous via tail-vein injection **(D)**, or intracranial **(F)** B16-F10 WT or NDUFS4 KO tumors. Significance determined by Log-rank test. **(C, E)** Tumor burden two weeks after B16-F10 WT or NDUFS4 KO tumors intravenous via tail-vein or intracranial injection, as measured by *ex vivo* bioluminescence of the lungs **(C)** or brain **(E)**, respectively. Y-axis represents the logarithmic-value of total flux. Significance was determined via two-sided Student’s *t*-test.

**Figure 5:**
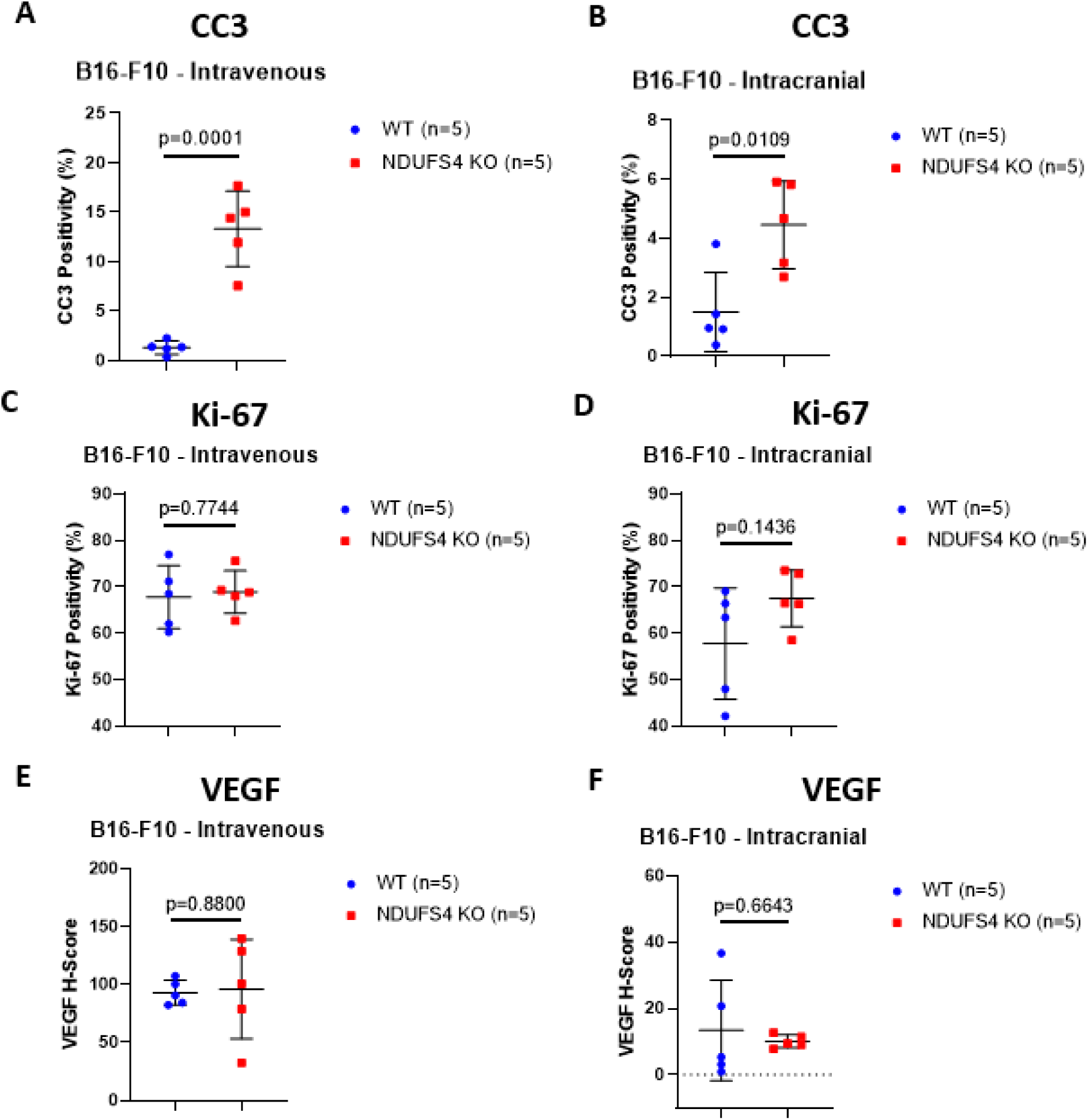
NDUFS4 KO results in increased apoptotic protein cleaved caspase 3 (CC3) compared to B16-F10 WT, but there is no impact on proliferation marker Ki-67 or angiogenic marker VEGF. Comparison of CC3 **(A-B)**, Ki-67 **(C-D)**, and VEGF **(E-F)** positivity between B16-F10 WT and NDUFS4 KO two weeks after intravenous injection and spread to the lungs **(A,C,E)** or direct intracranial injection **(B,D,F).** Y-axis represents staining positivity (CC3 and Ki-67) or H-Score (VEGF). Significance was determined via two-sided Student’s *t*-test.

## DISCUSSION

Developing an improved understanding of the predictors and drivers of distant metastasis remains a critical gap to improving outcomes in patients diagnosed with melanoma, the most aggressive of the common forms of skin cancer. Our findings here add to growing findings about the importance of metabolic pathways in this disease, and specifically implicate OXPHOS in the formation and growth of lung and brain metastases.

Using bulk gene expression analysis methods, we compared primary tumors from patients who developed metastases with primary tumors that failed to metastasize. OXPHOS gene expression was the most significantly enriched pathway between metastatic versus non-metastatic tumors. These findings were replicated in our transcriptomic analysis of genotype-matched metastatic versus non-metastatic murine samples. Collectively, these findings implicate OXPHOS as a pro-metastatic pathway. This conclusion is supported by the results seen in the RCAS-TVA mice, which is currently the only autochthonous melanoma model that accurately mimics the metastatic dynamics of melanoma and allows the assessment of candidate genes in this process.^[11]^ Genetic loss of PGC1α at tumor initiation did not impact primary tumor formation, but markedly inhibited metastatic spread to the lungs and brains. To further substantiate these findings, we demonstrated that suppression of the OXPHOS gene *Ndufs4* mitigated spread of B16-F10 and D4M-UV2 melanomas to the lungs and inhibited intracranial tumor growth. Cumulatively, these findings confirm that PGC1α/OXPHOS is necessary for lung and brain metastasis.

Metastases formed in the lungs of one mouse and lungs and brain of one mouse from the BCPP + myrAKT1 cohort. Interestingly, PCR analysis demonstrated incomplete loss of PGC1α in these mice. These findings serve as an internal control of the influence of PGC1α loss on metastasis formation within our BCPP RCAS-TVA model. The rarity of distant metastases in the BCPP + myrAKT1 RCAS-TVA model contrasts with the findings from our syngeneic models, which effectively bypass the process of primary tumor intravasation. Lung tumors formed in all mice with *Ndufs4* loss following intravenous injections, yet the overall tumor burden significantly decreased compared to the control group. Similarly, *Ndufs4* loss inhibited outgrowth of cells directly injected into the brains of mice without affecting the incidence of brain tumor formation. Collectively, these findings demonstrate that PGC1α/OXPHOS plays a critical role in intravasation, extravasation, and outgrowth of metastatic foci, with intravasation being particularly susceptible to loss of PGC1α/OXPHOS.

As PGC1α has been associated with activation of additional signaling pathways,^[18]^ our findings from the syngeneic models explicitly implicate OXPHOS as the means by which PGC1α facilitates metastasis. Interestingly, reduction in tumor burden observed in B16-F10 NDUFS4 KO tumors was not associated with changes in proliferation or angiogenesis, but with increased markers of apoptosis. Further studies are needed to better understand drivers of apoptosis in NDUFS4-deficient tumors and their impact on metastatic outgrowth in the lungs and brain.

Previously, we found that OXPHOS inhibitor treatment with IACS-010759 in BCP + myrAKT1 RCAS-TVA mice did not impact primary tumor growth or lung metastases, but completely inhibited brain metastasis formation.^[10]^ Our current findings contrast with these studies by demonstrating that suppression of PGC1α/OXPHOS at tumor initiation inhibits the general process of metastasis instead of being limited to only preventing brain metastasis formation. This discrepancy could be due to IACS-010759 treatment commencing after primary tumor formation in the prior studies. It is possible that cells had already seeded the lungs and brain and metastatic outgrowth was prevented in brain metastases to a more significant extent than in the lung given the increased OXPHOS reliance of tumors in the brain-metastatic niche.

Tumor cells experience a great deal of oxidative stress, particularly during metastasis, and decreased sensitivity to reactive oxygen species (ROS) is beneficial for melanoma cells. Our findings shed much needed light on the literature surrounding the role of PGC1α/OXPHOS in melanoma metastasis and how this signaling pathways fits in the greater schema of melanoma biology and response to ROS. PGC1α was first implicated in melanoma metastasis signaling as a mediator of resistance to targeted therapies.^[24, 25]^ Additional studies demonstrated that inhibition of PGC1α-mediated OXPHOS prevented metastatic spread of B16-F10 melanoma cells in a murine model of melanoma metastasis.^[26]^ Further studies showed that PGC1α overexpression promoted OXPHOS metabolism and significantly correlated with decreased overall survival in patients with advanced melanoma.^[27]^ Importantly, PGC1α loss inhibited expression of numerous ROS-scavenging genes and sensitized these cells to ROS. Later studies utilized a zebrafish model and demonstrated that PGC1α/OXPHOS promoted primary tumor progression and melanoma cell invasion and that PGC1α protein expression was positively associated with increased tumor thickness and expression of the proliferative marker Ki-67 and the reactive oxygen species SCARA3.^[28]^ Finally, others found that directly targeting OXPHOS triggered anoikis and impaired pulmonary metastasis in a B16-F10 melanoma model.^[29]^ Collectively, these findings demonstrate that PGC1α facilitates metastasis by inducing OXPHOS and mitigates potential detrimental side effects by activating antioxidant pathways.

Contrasting with these studies, Piskounova et al. demonstrated that melanoma cells respond to ROS by decreasing mitochondrial mass and mitochondrial membrane potential, suggesting that mitochondrial function is reduced in circulating melanoma cells.^[19]^ Importantly, comparisons in this study were made between distant metastases and circulating tumor cells and subcutaneous tumors. However, subcutaneous tumor growth is not synonymous with primary tumor development. It is entirely possible that while mitochondrial mass in distant metastases is lower than in subcutaneous tumors, the absolute level is still significantly higher than in true primary tumors. Luo et al. demonstrated that PGC1α expression inhibited invasive growth in local disease.^[18]^ When viewed in the context of their earlier work,^[27]^ the authors proposed a model in which melanomas are heterogeneous for PGC1α, and PGC1α-low cells are invasive while PGC1α-high cells are proliferative. However, our BCPP RCAS-TVA model contradicts this hypothesis. In our model, PGC1α is lost at the same time the melanocytes become oncogenic. We noted no difference in primary tumor incidence; in contrast, we observed a profound decrease in distant metastases due to loss of PGC1α. As previously mentioned, the only mice from the BCPP + myrAKT1 cohort that developed metastases had incomplete loss of PGC1α. The discrepancies between our data and Luo et al. likely stem from inherent differences in our models, as their studies were performed predominantly in manipulated human melanoma cell lines *in vitro* and *in vivo*. Similar to Piskounova et al, *in vivo* experiments by Luo et al. featured frequent comparisons between distant metastases, circulating tumor cells, and subcutaneous tumors as opposed to comparisons with autochthonous primary tumors. While their microarray studies demonstrated that PGC1α decreased as melanomas transitioned from radial to vertical growth phases, Salhi et al. demonstrated the opposite findings using immunohistochemistry.^[28]^ Our transcriptomic studies of murine and human samples demonstrated that OXPHOS gene expression in primary tumors predicted increased distant metastasis risk. Based on *in vitro* analysis, we did not observe changes in cytoplasmic or mitochondrial ROS levels in B16-F10 or D4M-UV2 cells compared to NDUFS4 KO cells despite increased OXPHOS. However, we identified enrichment of several ROS signaling pathways in metastatic versus non-metastatic primary tumors from both mice and patients.

Maintenance of PGC1α expression thus affords metastatic melanoma cells two benefits in their spread to distant organs. First, these cells are better equipped to handle oxidative stress secondary to interactions with their microenvironment or intrinsically from OXPHOS. Second, the OXPHOS pathway itself affords the metastatic cells decreased reliance on glucose and increased ability to scavenge foreign soils for a wider range of metabolic fuels, including fatty acids, acetate, branched-chain amino acids, and glutamine.^[30–32]^ Our findings could lay the groundwork for the development of strategies aimed at treating and preventing metastasis from advanced melanoma by targeting tumor OXPHOS or the energetic dependencies inherent in these tumors.

In summary, our clinical cohort of primary melanomas, an autochthonous RCAS-TVA model, and syngeneic models demonstrate that tumor PGC1α/OXPHOS plays a crucial role in the formation and growth of melanoma metastases in the lungs and brain. These experiments provide a foundation for investigating the role of tumor OXPHOS in melanoma metastases to other organs as well as in distant metastases from other tumor types. This improved understanding of melanoma metabolism takes a critical step forward in identifying novel vulnerabilities for surveillance/preventative strategies for metastasis.

## METHODS

### Patient Samples

Sample selection and acquisition have been previously described for the primary tumor cohort.^[21]^ Briefly, patients between 1998-2010 were included if they had a single, invasive, primary melanoma with Breslow thickness >1.5mm, FFPE primary tumor tissue available for analysis, no clinical evidence of regional metastasis at the time of primary tumor diagnosis, and a sentinel lymph node biopsy conducted <1 year post-diagnosis.

### Cell Lines

All mammalian cell lines were grown at 37°C under 5% CO_2_. B16-F10/B16-F10 NDUFS4 KO cells [provided by Gregory Delgoffe, UPMC] and D4M-UV2 cells [provided by David Fisher, MGH]/NDUFS4 KO cells [developed/validated by Synthego], were grown in DMEM media supplemented with glutamine, 10% Non-Essential Amino Acids (NEAA), and 10% heat-inactivated FBS (all Corning, Inc.). DF-1 avian fibroblasts were grown in DMEM-high glucose media (Thermo Fisher) supplemented with 10% FBS (Atlanta Biologicals) and 0.5 μg/mL Gentamicin (Thermo Fisher), and maintained at 39°C. All cell lines were confirmed mycoplasma-negative using the MycoAlert Mycoplasma Detection Kit (Lonza).

### Mice

All mouse experiments were approved by the Institutional Animal Care and Use Committees of University of Utah Health Sciences Center and MDACC. All experiments using the *Dct::TVA;BRAF^CA^;Cdkn2a^lox/lox^:Pten^lox/lox^* (BCP) and *Dct::TVA;BRAF^CA^; Cdkn2a^lox/lox^:Pten^lox/lox^;Pgc1α^lox/lox^* (BCPP) RCAS-TVA mice were conducted at the University of Utah Health Sciences Center. DNA from tail biopsies was used to genotype the *TVA* transgene, *BRAF^CA^, Cdkn2a^lox^, Pten^lox^*, and wild-type alleles as described.^[33–35]^ ‘PCR to detect a 420-bp *Pgc1α^lox^* allele and a 360 bp wild-type allele was carried out with the following primer sequences: Fwd 5’-TCCAGTAGGCAGAGATTTATGAC-3’; Rev 5’-TGTCTGGTTTGACAATCTGCTAGGTC-3’. Experiments using C57BL/6J mice (female; 6-8 weeks; Jackson) were performed at the MDACC South Campus Animal Vivarium.

### RNA-sequencing Analyses

Raw counts files were acquired, and the OXPHOS-Index (OP-Index) was derived, as previously described.^[10]^ Briefly, comparisons of interest were performed using functions from the *edgeR* and *limma/voom* Bioconductor packages in R (v3.6.1)(8). Preranked GSEA (GSEA-P) was implemented using the GenePattern module *GSEAPreranked* (v6.0.10). Rank metric was calculated as the sign of log2-FCs multiplied by the inverse of *p*-values calculated using the EdgeR/limma/voom pipeline. ssGSEA was conducted on TMM-normalized, voom-transformed log2-(CPM+0.5) expression matrices using the GenePattern module *ssGSEAProjection* (v9.0.10) settings to calculate enrichment scores for the 8 OXPHOS-related gene sets.^[10]^

### RCAS-TVA Model Tumor Induction and Sample Collection

#### Tumor induction

Cloning details for RCAS-AKT mutant constructs, method of viral propagation, and generation of primary tumors in BCP mice using RCAS-*Cre* ± RCAS-*Akt1/2/*3^E17K^ mutant retroviruses have been described.^[20]^ Primary tumors were generated in *Dct::TVA;Braf^CA^*; *Cdkn2a^lox/lox^*;*Pten^lox/lox^* (BCP) and *Dct::TVA;BRAF^CA^; Cdkn2a^lox/lox^:Pten^lox/lox^*;*Pgc1a^lox/lox^*(BCPP) mice using RCAS-*Cre* ± RCAS-myr*Akt1* retroviruses as previously described.^[11]^

#### FFPE specimens

A full necropsy was performed on all mice following euthanasia. Brain, lung, and primary tumor tissues were fixed in formalin overnight, dehydrated in 70% ethyl alcohol, processed, and paraffin-embedded. Sections were stained with hematoxylin & eosin (H&E) for review by a pathologist.

#### Frozen primary tumor specimens

Frozen primary tumor samples were embedded in optimal cutting temperature compound (OCT) and brought to −20°C. A 5 µm H&E-stained slide was prepared from each sample using a cryostat and reviewed by a pathologist for regions containing 70% or more viable tumor. Marked H&E slides were then used to guide macrodissection of viable tumor from the OCT blocks. For RNA extraction and DNase treatment, the Roche High Pure miRNA Isolation Kit was used as previously described.^[20]^ To confirm that the PGC1α allele was fully recombined in all tumors, DNA was isolated from frozen primary tumors and the PGC1α recombined allele (expected band size: 420 bp) was amplified by PCR with the following primer sequences: 1) 5’-TCCAGTAGGCAGAGATTTATGAC-3’ and 2) 5’-CCAACTGTCTATAATTCCAGTTC-3’.

#### CRISPR *Ndufs4* knockout syngeneic injection models

1×10^6^ B16-F10 WT/NDUFS4 KO or D4M-UV2 WT/NDUFS4 KO cells suspended in HBSS were injected subcutaneously. Tumors were measured using digital calipers every three days. 1×10^6^ B16-F10 WT/NDUFS4 KO or D4M-UV2 WT/NDUFS4 KO cells were injected intravenously in mice. 5×10^3^ B16-F10 WT/NDUFS4 KO or D4M-UV2 WT/NDUFS4 KO cells were directly implanted in the brain parenchyma of mice. Mice were weighed every 3 days. Mice that developed >20% weight loss, neurological symptoms, or respiratory distress were euthanized. Additional mice were injected intravenously or intracranially and harvested at two weeks (B16-F10/NDUFS4 KO) or at three weeks (D4M-UV2/NDUFS4 KO) and tumor burden of the lungs or brain was measured via *ex vivo* bioluminescence (Supplemental Methods).

#### *CRISPR* Ndufs4 *knockout syngeneic injection models*

1×10^6^ B16-F10 WT/NDUFS4 KO or D4M-UV2 WT/NDUFS4 KO cells suspended in HBSS were injected subcutaneously. Tumors were measured using digital calipers every three days. 1×10^6^ B16-F10 WT/NDUFS4 KO or D4M-UV2 WT/NDUFS4 KO cells were injected intravenously in mice. 5×10^3^ B16-F10 WT/NDUFS4 KO or D4M-UV2 WT/NDUFS4 KO cells were directly implanted in the brain parenchyma of mice. Mice were weighed every 3 days. Mice that developed >20% weight loss, neurological symptoms, or respiratory distress were euthanized. Additional mice were injected intravenously or intracranially and harvested 2 weeks later (B16-F10/NDUFS4 KO) or two weeks later (D4M-UV2/NDUFS4 KO) and tumor burden of the lungs or brain was measured via *ex vivo* bioluminescence (Supplemental Methods).

### Statistical Analyses

Overall survival was defined as the time interval from date of tumor injection to date of harvest. Survival duration was analyzed by the Kaplan-Meier method. Survival curves were drawn in Prism 9.0 (Graphpad). Hazard ratios and significance were calculated via the Mantel-Haenszel test and log-rank test, respectively. Additional data analyses and representations were performed either with the R (v3.6.1), Microsoft Excel, or Prism 9.0 (GraphPad). Comparison of continuous variables between two groups was performed by unpaired Student’s *t*-test. Evaluation of associations between categorical variables was evaluated by Fisher’s exact test. Lastly, all statistical significance testing was two-sided at Type-I error rate of 0.05.

## Author Contributions

Conception and design: RAG, GMF, DAK, MAD, SLH; Acquisition of data: RAG, GMF, DAK, JRC, CWH; Analysis and interpretation of data: RAG, GMF, DAK, AYJ, DAL, AHG, RNLS, SLH, MAD; Writing, review, and/or revision of the manuscript: All authors; Study supervision: RAG, GMF, DAK, SLH, MAD

