## Supplement Tables for "Oxidative Phosphorylation (OXPHOS) Promotes the Formation and Growth of Melanoma Lung and Brain Metastases"

**Supplemental Tables**

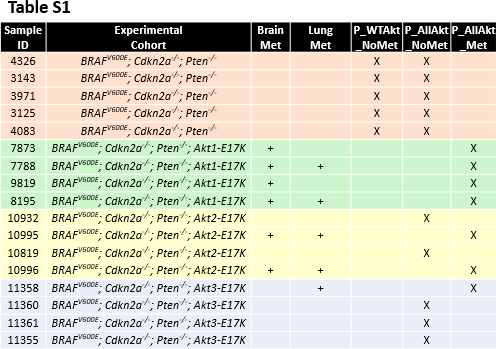

**Supplemental Table 1:** Genotypes and metastasis incidence of primary tumors acquired from the BCP ± AKT1/2/3-E17K RCAS-TVA model of spontaneous brain and lung metastasis. “Sample ID” is the ear tag of the mouse. “Experimental Cohort” is the genotype of each mouse’s tumor. “Brain Met” and “Lung Met” indicate the presence (+) or absence (blank) of metastases at the designated anatomical site. “P_WTAkt_NoMet”, “P_AllAkt_NoMet”, and “P_AllAkt_Met” indicate GSEA-P analysis groups that samples were assigned to (X).

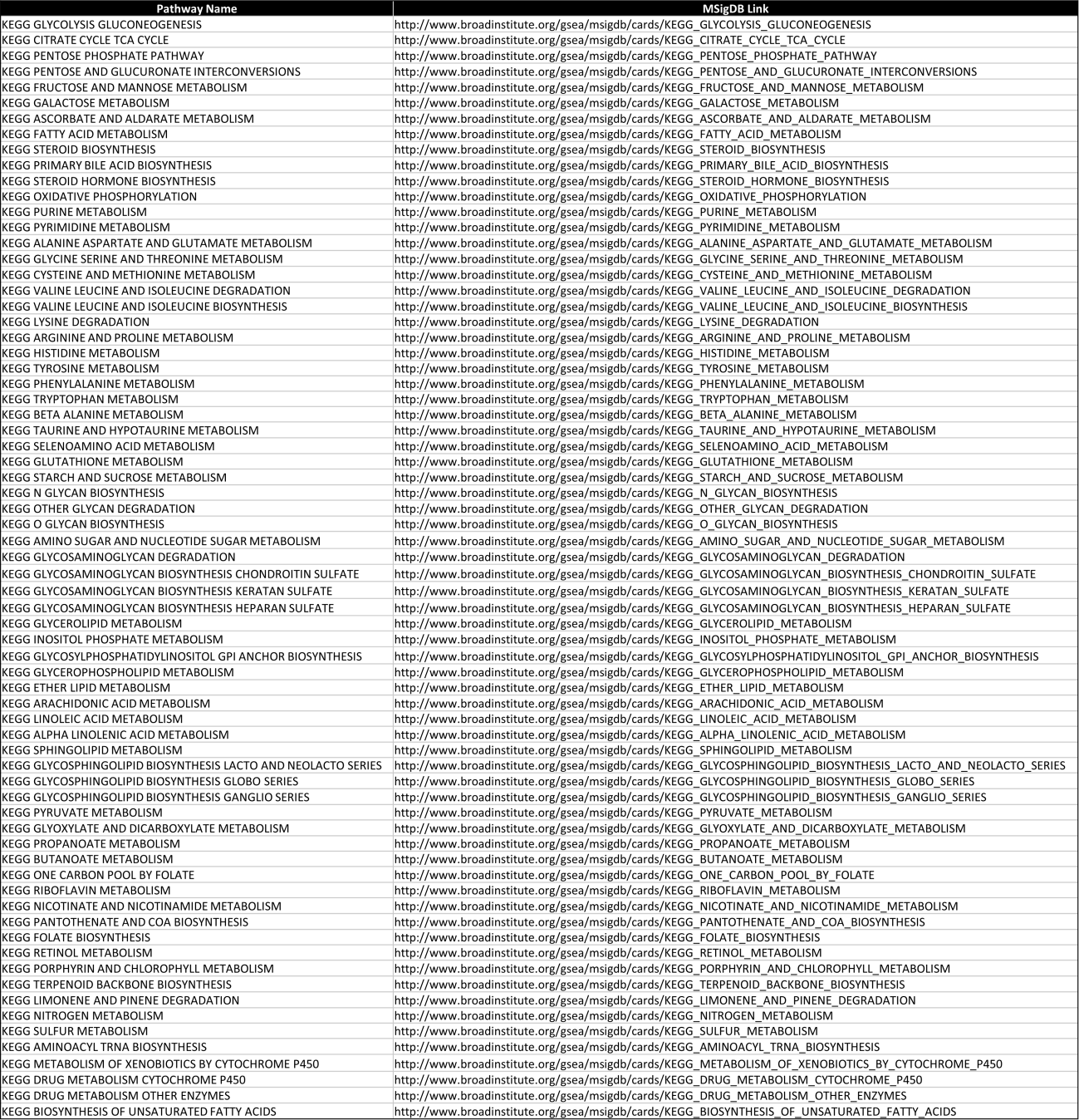

**Supplemental Table 2:** KEGG metabolism gene sets used in metabolomics pre-ranked gene set enrichment (GSEA-P) analysis.

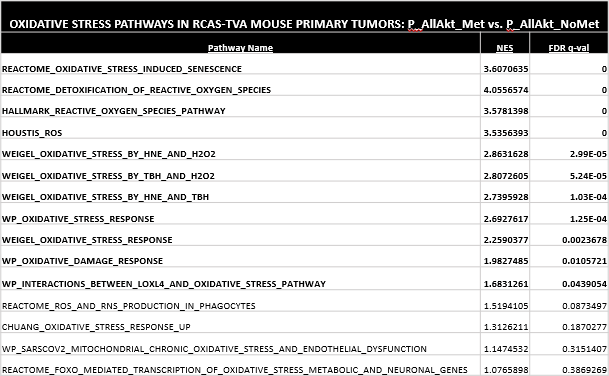

**Supplemental Table 3:** Oxidative stress pathways curated from Hallmark, C1, and C2 gene sets that are enriched (FDR q-val <0.05) in P_AllAkt_Met versus P_AllAkt_NoMet primary tumors in BCP AKT1/2/3-E17K RCAS-TVA mice.

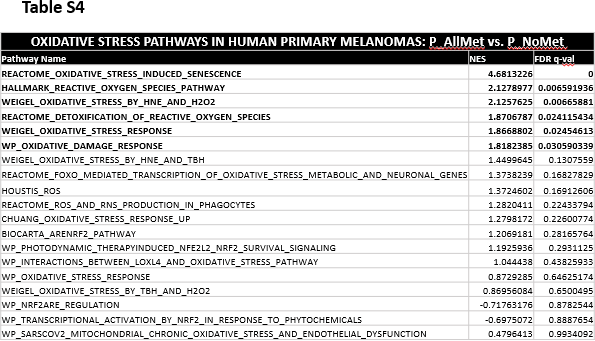

**Supplemental Table 4:** Oxidative stress pathways curated from Hallmark, C1, and C2 gene sets that are enriched (FDR q-val <0.05) in P_AllMet versus P_NoMet primary tumors in melanoma patients.
