## Supplement Methods for "Oxidative Phosphorylation (OXPHOS) Promotes the Formation and Growth of Melanoma Lung and Brain Metastases"

**Supplemental Methods Bioenergetics Stress Test**

A Seahorse XFe96 Bioanalyzer (Agilent) was used according to the manufacturer’s instructions. Briefly, D4M-UV2/*Ndufs4*-deficient D4M-UV2 or B16-F10/*Ndufs4*-deficient B16-F10 cells were plated in a 96-well Seahorse XF Cell Culture Microplate at a density of 25,000/well in 100 ul of the appropriate cell culture growth media and incubated for 16 hours at 37°C under 5% CO_2_. MitoStress Test was then performed as previously described.^[10]^

**Mitochondrial Flow Cytometry Assays (ROS, Mitochondrial Mass, Mitochondrial Membrane Potential)**

We stained cells for cytoplasmic ROS (CellROX DeepRed; 5μM) or mitochondrial ROS (CellROX Green; 5μM) (all Life Technologies) for 30 minutes at 37°C under 5% CO_2_ and then stained with Live/Dead SYTOX Blue (5μM), as previously described.^[19]^ Cells were analyzed immediately using a Fortessa X-20 flow cytometer and analyzed using FlowJo 10.8.1.

***Ex Vivo* Bioluminescence**

Mice were injected intraperitoneally with D-luciferin (150mg/kg; BioVision), sacrificed 5 minutes later, brain/lungs harvested, and imaged on an IVIS100 imager at 10 minutes post-luciferin injection. Signal intensities were quantified as total flux using Living Image (v4.5.2) software.

**Immunohistochemistry**

All IHC studies were performed on 4 μm FFPE sections using a Leica BOND RXm autostainer. Slides were stained with antibodies targeting Cleaved Caspase 3 (CC3;

Clone D3E9; Dilution 1:250; Cell Signaling #9579), Ki-67 (Clone D3B5; Dilution 1:400; Cell Signaling #12202), and VEGFA (polyclonal; Dilution 1:200; Atlas Antibodies #HPA069116) using a modified version of the standard Leica Bond DAB “F” IHC protocol. Image analysis was performed using Aperio automated algorithms as previously described.^[10]^ CC3 was scored as % of tumor cells showing cytoplasmic and perinuclear positivity. Ki-67 was scored as % of tumor cells showing nuclear positivity. H-scores for VEGF was scored based on % of tumor cells showing membranous and cytoplasmic positivity and intensity of staining. Significance determined via two-sided Student’s *t*-test.
