## Supplement Figures for "Oxidative Phosphorylation (OXPHOS) Promotes the Formation and Growth of Melanoma Lung and Brain Metastases"

**Supplemental Figures**

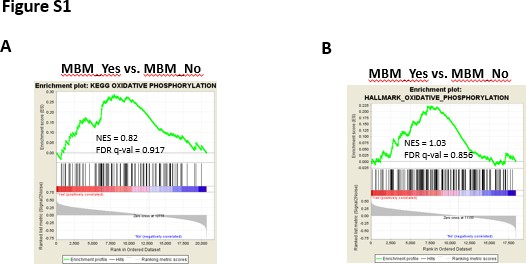

**Supplement	Figure	1:	Oxidative	phosphorylation	does	not	correlate	with cerebrotropism in melanoma extracranial metastases.** GSEA enrichments plots of

**(A)** KEGG OXPHOS and (**B)** MSigDB Hallmarks OXPHOS gene sets in extracranial metastases with cerebrotropism (development of brain metastasis with <6 months of stage IV disease diagnosis) vs. extracranial metastases that failed to develop brain metastases for >18 months from stage IV diagnosis. Normalized enrichment score (NES) and FDR q-val are listed on the enrichment plots.

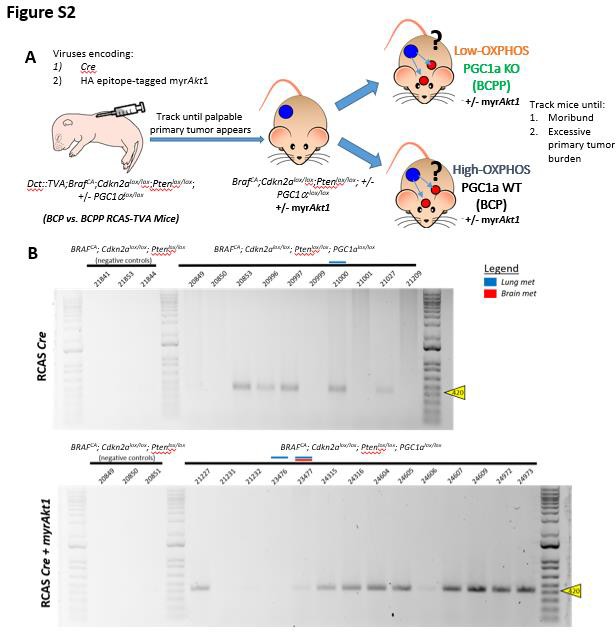

**Supplemental Figure 2: Allelic deletion of PGC1α used to evaluate the role of OXPHOS in an RCAS-TVA model of spontaneous murine melanoma brain and lung metastasis. (A)** Experimental design used to functionally validate the role of OXPHOS in brain and lung metastasis from primary melanoma tumors in the BCP vs. BCPP RCAS-TVA models. **(B)** PCR gel correlating to tumors from BCPP RCAS-TVA mice evaluating loss of *PGC1α*. Numbers indicate mouse IDs. Blue and red marks

indicate BCPP RCAS-TVA mice that developed spontaneous lung and brain metastases, respectively. Yellow triangles mark distance on PCR gels that correlate with distance of PGC1α bands.

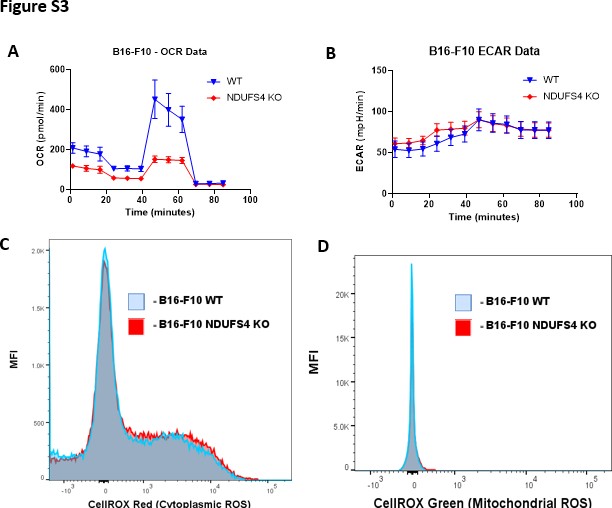

**Supplemental Figure 3: NDUFS4 KO reduces oxidative phosphorylation (OXPHOS) while maintaining stable glycolysis and ROS compared to B16-F10 WT. (A-B)** Seahorse mitochondrial stress test was performed on B16-F10 WT and NDUFS4 KO cells grown in vitro. The Y-axis measures oxygen consumption rate (OCR; **A**) and extracellular acidification rate (ECAR; **B**), which are measures of OXPHOS and glycolysis, respectively. The X-axis designates timepoints as cells undergo 3 measurements of basal, oligomycin-inhibited, FCCP-inhibited, and rotenone/antimycin A-inhibited OCR each. **(C-D)** Flow cytometry histograms analyzing fluorescent staining of cytoplasmic ROS via CellROX Red **(C)** and mitochondrial ROS via CellROX Green

**(D)** in B16-F10 WT and NDUFS4 tumor cells grown in vitro. The Y-axis represents median fluorescence intensity (MFI).

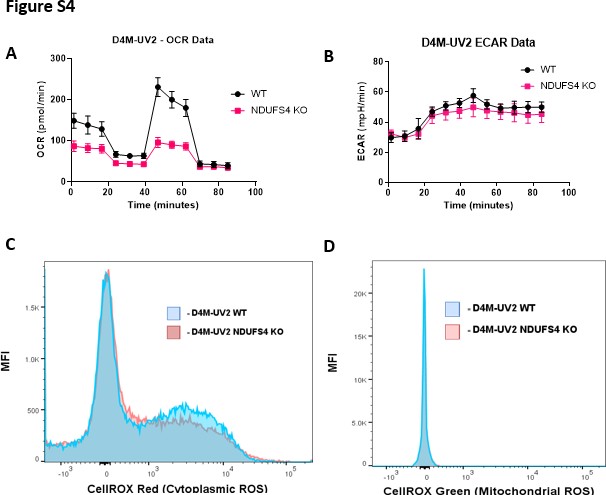

**Supplemental Figure 4: NDUFS4 KO reduces oxidative phosphorylation (OXPHOS) while maintaining stable glycolysis and ROS compared to D4M-UV2 WT. (A-B)** Seahorse mitochondrial stress test was performed on D4M-UV2 WT and NDUFS4 KO cells grown in vitro. The Y-axis measures oxygen consumption rate (OCR; **A**) and extracellular acidification rate (ECAR; **B**), which are measures of OXPHOS and glycolysis, respectively. The X-axis designates timepoints as cells undergo 3 measurements of basal, oligomycin-inhibited, FCCP-inhibited, and rotenone/antimycin A-inhibited OCR each. **(C-D)** Flow cytometry histograms analyzing fluorescent staining of cytoplasmic ROS via CellROX Red **(C)** and mitochondrial ROS via CellROX Green

**(D)** in D4M-UV2 WT and NDUFS4 tumor cells grown in vitro. The Y-axis represents median fluorescence intensity (MFI).

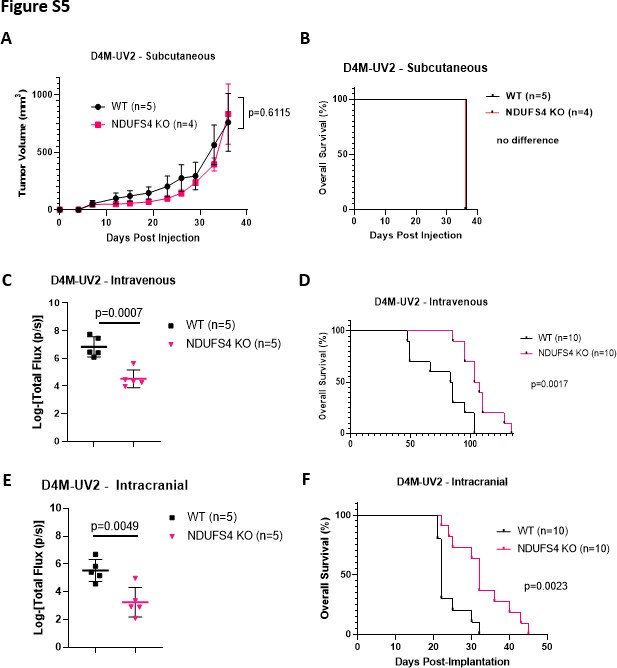

**Supplemental Figure 5: NDUFS4/OXPHOS is functionally significant for D4M-UV2 melanoma tumor growth and overall survival in syngeneic injection models of lung and brain, but not subcutaneous, tumor growth. (A)** Growth of subcutaneous tumors. **(B, D, F)** Kaplan-Meier analysis of overall survival for mice bearing

subcutaneous **(B)**, intravenous via tail-vein injection **(D)**, or intracranial **(F)** D4M-UV2

WT or NDUFS4 KO tumors. Significance determined by Log-rank test. **(C, E)** Tumor burden two weeks after D4M-UV2 WT or NDUFS4 KO tumors intravenous via tail-vein or intracranial injection, as measured by *ex vivo* bioluminescence of the lungs **(C)** or brain **(E)**, respectively. Y-axis represents the logarithmic-value of total flux. Significance was determined via two-sided Student’s *t*-test.
